# Brain tissue cerebrospinal fluid fraction increases quadratically in normal aging

**DOI:** 10.1101/2023.02.09.527912

**Authors:** Liangdong Zhou, Yi Li, Elizabeth M. Sweeney, Xiuyuan H. Wang, Amy Kuceyeski, Gloria C. Chiang, Jana Ivanidze, Yi Wang, Susan A. Gauthier, Mony J. de Leon, Thanh D. Nguyen

**Affiliations:** Department of Radiology, Weill Cornell Medicine, New York, NY, USA; Penn Statistics in Imaging and Visualization Endeavor (PennSIVE), Department of Biostatistics and Epidemiology, University of Pennsylvania, Philadelphia, USA; Department of Statistics and Data Science, Cornell University, Ithaca, NY, USA

**Keywords:** T2 relaxometry, cerebrospinal fluid fraction (CSFF), myelin water fraction (MWF), intra/extra-cellular water fraction (IEWF), FAST-T2, normal aging, perivascular space

## Abstract

**Background and Purpose:** Our objective was to apply multi-compartment T2 relaxometry in cognitively normal individuals aged 20-80 years to study the effect of aging on the parenchymal cerebrospinal fluid fraction (CSFF), a measure of the microscopic-scale CSF space.

**Materials and Methods:** A total of 66 volunteers (age range, 22-80 years) were enrolled. Voxel-wise maps of short-T2 myelin water fraction (MWF), intermediate-T2 intra/extra-cellular water fraction (IEWF), and long-T2 CSFF were obtained using fast acquisition with spiral trajectory and adiabatic T2prep (FAST-T2) sequence and three-pool non-linear least squares fitting.

Multiple linear regression analysis with correction for multiple comparisons was performed to study the association between age and regional MWF, IEWF, and CSFF measurements, adjusting for sex and region of interest (ROI) volume. The cerebral white matter (WM), cerebral cortex, and subcortical deep gray matter (GM) were considered as ROIs. In each model, a quadratic term for age was tested using an ANOVA test. A Spearman’s correlation between the normalized lateral ventricle volume, a measure of organ-level CSF space, and the regional CSFF, a measure of tissue-level CSF space, was computed.

**Results:** In the multiple regression analysis, we found a statistically significant quadratic relationship between age and regional CSFF for all three ROIs (all p-values < 0.001). A statistically significant quadratic relationship with age was also found for MWF in the deep GM (p = 0.004) and IEWF in the cortex (p = 0.012). There was a statistically significant linear relationship between age and regional IEWF in the cerebral WM (p = 0.006) and deep GM (p = 0.002). In the univariate correlation analysis, the normalized lateral ventricle volume was found to correlate moderately with the regional CSFF measurement in the cerebral WM (ρ = 0.43, p < 0.001), cortex (ρ = 0.43, p < 0.001), and deep GM (ρ = 0.49, p < 0.001).

**Conclusion:** Brain tissue water residing in different water compartments shows complex changing patterns with age. Parenchymal CSFF, a biomarker of microscopic-scale CSF-like water, shows a quadratic increase in both GM and WM, starting approximately at the age of 50.

## INTRODUCTION

The cerebrospinal fluid (CSF) system plays a prominent role in maintaining the homeostasis of the central nervous system, providing hydromechanical protection, nutrient transport, and metabolic waste removal, among other essential functions (Sakka et al., 2011). In the brain, CSF occupies the ventricles, the cranial subarachnoid space, and the perivascular space (PVS). PVS surrounds blood vessels and acts as a conduit for the exchange between CSF and the interstitial fluid (Wardlaw et al., 2020). Advanced imaging techniques have shown that PVS is a key component of the glymphatic system for brain waste removal (Weller et al., 1992;Iliff et al., 2012;Nedergaard, 2013;Taoka et al., 2017). PVS dilation occurs in normal aging (Kim et al., 2022) and has been established as an early imaging marker of cerebral small vessel disease (Wardlaw et al., 2013), cerebral amyloid angiopathy (van Veluw et al., 2016;Charidimou et al., 2017), and Alzheimer’s disease (Banerjee et al., 2017;Boespflug et al., 2018). Traditionally, visible PVS enlargement on T2-weighted (T2W) image is assessed by counting or segmenting hyperintense punctate foci in the white matter (WM) centrum semiovale and the basal ganglia (Potter et al., 2015;Ramirez et al., 2016;Ballerini et al., 2018). However, this method cannot reliably detect microscopic-scale PVS which is much smaller than the typical 1 mm resolution of conventional MRI.

Multi-component T2 relaxometry is a quantitative MRI method that can separate signals of different water compartments within an imaging voxel based on their T2 relaxation times (Whittall et al., 1997). In healthy brain gray matter (GM) and WM tissues, three water compartments can be distinguished based on the degree of water mobility, including water trapped in the myelin sheath with short T2 (∼10 ms), intra/extra-cellular water with intermediate T2 (∼70 ms) such as axonal and interstitial water, and CSF with long T2 (∼2 s) (MacKay et al., 2006). Of these, myelin water fraction (MWF), expressed as the ratio between the myelin water volume and the total water volume in a voxel (MacKay et al., 1994), has been established as an imaging marker of myelin injury in multiple sclerosis (MS) and other demyelinating disorders (Laule et al., 2008;Mancini et al., 2020;van der Weijden et al., 2021). This has spurred the development of a number of fast MRI techniques for MWF mapping (Alonso-Ortiz et al., 2015), including 3D gradient echo spin echo (GRASE) (Prasloski et al., 2012), 3D multi-component driven equilibrium steady-state observation of T1 and T2 (mcDESPOT) (Deoni et al., 2008), and 3D fast acquisition with spiral trajectory and adiabatic T2prep (FAST-T2) (Nguyen et al., 2016) sequences. To date, however, there have been few MRI studies on the effect of age on parenchymal CSF fraction (CSFF), defined as the ratio between the CSF volume and the total water volume in a voxel and could serve as a quantitative biomarker of PVS dilation on the microscopic scale. In a recent study (Canales-Rodriguez et al., 2021), Canales-Rodríguez et al applied multi-component T2 relaxometry to the brain data of cognitively normal (CN) subjects obtained with the GRASE sequence and reported a linear increase of CSFF with age. However, their imaging voxel size was rather large (42.9 mm^3^) and elderly subjects aged 60 years or above were not included. The objective of this study was to demonstrate the feasibility of the FAST-T2 sequence with a smaller voxel size (7.8 mm^3^) for CSFF mapping and to study the effect of aging on parenchymal CSFF in CN individuals aged 20-80 years.

## METHODS

### Study participants

This was a cross-sectional study conducted in a cohort of 66 CN volunteers (26 men (39.4%), 40 women (60.6%); mean age, 48.4 years ±16.7 (standard deviation [SD]); age range, 22-80 years) who had brain MRI with FAST-T2 sequence performed as part of imaging research studies at Weill Cornell Medicine. All studies were approved by the local institutional review board and written informed consent was obtained from all participants prior to imaging. Subjects contraindicated for MRI, those with a history of neurological problems (including, but not limited to, stroke, head trauma, brain infarcts, intracranial mass, MS, meningitis, encephalitis, prior brain surgery), or those on medications that may affect cognitive performance, were excluded. Twenty-three out of 30 subjects aged 50 years or more also underwent extensive neurocognitive assessments which included the Brief Cognitive Rating Scale (Reisberg and Ferris, 1988), the Global Deterioration Scale (GDS) (Reisberg et al., 1982), and the Clinical Dementia Rating (CDR) (Morris, 1993).

### MRI examination

All 66 subjects underwent brain MRI examination on Siemens 3T scanners (Siemens Healthineers, Erlangen, Germany). Of these, 20 subjects (30.3%) were scanned on Skyra scanners using a product 20-channel head/neck coil and 46 subjects (69.7%) were scanned on a Prisma scanner using a product 64-channel head/neck coil. The brain imaging protocol consisted of 3D T1-weighted (T1W) magnetization-prepared rapid acquisition gradient echo (MPRAGE) and 2D T2W turbo spin echo (TSE) or 3D T2W Sampling Perfection with Application optimized Contrast using different flip angle Evolution (SPACE) sequences for anatomical structure, as well as 3D FAST-T2 sequence for water fraction (WF) mapping, among others. The imaging parameters were as follows: 1) 3D sagittal MPRAGE: TR/TE/TI = 2300/2.3/900 ms, flip angle (FA) = 8°, readout bandwidth (rBW) = 200 Hz/pixel, voxel size = 1.0 mm isotropic, GRAPPA parallel imaging factor (R) = 2, scan time = 5.5 min; 2a) 2D axial T2W TSE: TR/TE = 5840/93 ms, FA = 90°, rBW = 223 Hz/pixel, turbo factor = 18, number of signal averages (NSA) = 2, voxel size = 0.5×0.8×3.0 mm^3^, R = 2, scan time = 4 min; or alternatively, 2b) 3D sagittal T2W SPACE: TR/TE=3200/408 ms, FA = 90°, rBW = 751 Hz/pixel, turbo factor = 285, voxel size = 1.0 mm isotropic; 3) 3D axial FAST-T2 (Nguyen et al., 2016): spiral TR/TE = 7.8/0.5 ms, nominal T2prep times = 0 (T2-prep turned off), 7.5, 17.5, 67.5, 147.5, and 307.5 ms, T1 saturation recovery time (wait time between the saturation pulse at the end of spiral readout and the next T2prep) = 2 s, FA = 10°, rBW = 1042 Hz/pixel, number of spiral leaves per stack = 32, number of spiral leaves collected per T2prep = 64, voxel size = 1.3×1.3×5 mm^3^, scan time = 4 min.

### Image post-processing

The acquired images were processed using an automated pipeline based on FMRIB Software Library (FSL) v6.0 (Jenkinson et al., 2012). This consisted of applying FAST bias field correction (Zhang et al., 2001) and BET brain extraction (Smith, 2002) to the structural T1W image, followed by rigid-body registration (six degrees of freedom) of the T1W brain mask to the FAST-T2 image using FLIRT linear registration algorithm (Jenkinson and Smith, 2001). In addition, regional brain segmentation and cortical parcellation (aparc+aseg) was obtained from T1W and T2W images using FreeSurfer v7.1 recon-all command (Fischl, 2012), from which the binary tissue mask for our three regions of interest (ROIs), including the cerebral WM, the cerebral cortex, and the subcortical deep GM (cerebral GM minus cerebral cortex), were derived. To reduce the influence of CSF occupying the brain ventricles and the subarachnoid space on regional CSFF measurements in these ROIs due to the partial volume effect at the tissue-CSF interface, more conservative tissue ROIs were generated for regional WF measurements by first applying a 1 mm isotropic erosion to the original FreeSurfer ROIs, followed by another erosion such that the distance between the ROI boundary and the boundary of these CSF-filled spaces was at least 1 mm in-plane and 5 mm through-plane (chosen based on the 1.3×1.3×5 mm^3^ voxel size of the FAST-T2 acquisition).

To mitigate Gibbs ringing artifacts present in the FAST-T2 images (predominantly in the slice direction due to the relatively thick 5 mm acquired slices), a local subvoxel-shifts Gibbs correction algorithm (Kellner et al., 2016) was applied (mrdegibbs command in MRtrix3 software package (Tournier et al., 2019)). Additionally, we applied correction for the noise bias inherent in the FAST-T2 sum-of-squares coil-combined magnitude data, which follows noncentral chi distribution (Constantinides et al., 1997) and mainly affects the later low signal-to-noise ratio (SNR) echoes (the tail of the T2 signal decay curve). For this purpose, the number of the coil elements active during the FAST-T2 acquisition was determined from the DICOM image header, and the magnitude noise statistics was estimated from ROIs placed in the background air region of the two edge slices acquired at the last TE (these were chosen to minimize the effect of the anatomical signal for accurate noise estimation). The corrected FAST-T2 signal amplitude was then obtained from the measured noisy magnitude signal on a per-voxel basis by using a lookup table calculated for the given number of active coil elements following Eq.2 in (Constantinides et al., 1997).

Whole-brain MWF, intra/extra-cellular water fraction (IEWF), and CSFF maps were computed from the Gibbs- and noise-corrected FAST-T2 magnitude image data using a multi-voxel three-pool nonlinear least-squares fitting algorithm with a Laplacian local spatial smoothness constraint (Andrews et al., 2005;Nguyen et al., 2016). The lower and upper T2 bounds for each of the three water pools (in milliseconds) were set to [5 20], [20 200], and [200 2000] as in (Nguyen et al., 2016). Following the conventional T2 relaxometry nomenclature (Lancaster et al., 2003;MacKay et al., 2006;Canales-Rodriguez et al., 2021), these water compartments represent short-T2 myelin water, intermediate-T2 intra/extra-cellular water (e.g., axonal and interstitial water), and long-T2 free and quasi-free water (mainly CSF) (MacKay et al., 2006), respectively. At each voxel, the T2 value of the myelin water and CSF components was initialized to 10 ms and 2000 ms (Spijkerman et al., 2018), respectively, while the initial T2 value of the intra/extra-cellular water component was calculated using an efficient linear least-squares fit of the natural logarithm of the first four echoes (Otto et al., 2011). A limited-memory Broyden-Fletcher-Goldfarb-Shanno (L-BFGS) algorithm implemented in MATLAB R2022a (The MathWorks Inc, Natick, MA, USA) was used to obtain the constrained nonlinear least-squares solution, resulting in three compartmental water T2 values and three corresponding water signal amplitudes per image voxel (Schmidt et al.). The relative MWF, IEWF, and CSFF maps were calculated as the ratio of the signal of the myelin water, intra/extra-cellular water, and CSF, respectively, and the total water signal within a voxel. The water maps were then aligned to the FreeSurfer space by rigid-body registration (six degrees of freedom) using the FLIRT algorithm.

### Statistical analysis

The statistical analysis was performed in the R environment (version 4.2.2; R Foundation for Statistical Computing, Vienna, Austria). Data are presented as mean ± SD. A p-value of less than 0.05 was considered statistically significant. Following visual inspection of the extracted mean MWF, IEWF, and CSFF data obtained in the cerebral deep GM, cerebral cortex, and cerebral WM as a function of age, we considered two multiple linear regression models with each of these regional WF measurements as an outcome variable (9 in total) and age as the main predictor variable of interest, adjusting for sex and ROI volume (uneroded and normalized to the subject skull size using SIENAX algorithm (Smith et al., 2002) to account for the effect of brain atrophy in normal aging (Resnick et al., 2003;Raz et al., 2004)):

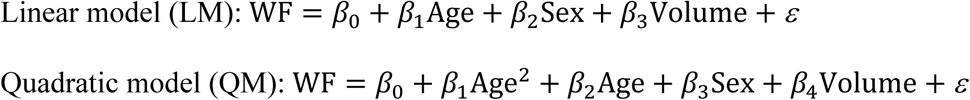

The final regression model was selected as the most parsimonious model that still fits the data adequately. For this purpose, an ANOVA test was performed to test if the squared term for age should be included in the model. If the difference was statistically significant, QM was selected, otherwise LM was selected. To correct for multiple comparisons, the p-values obtained for the age variable from the 9 regression models were adjusted using the false discovery rate (FDR) method (Benjamini and Hochberg, 1995). Finally, to assess the relationship between CSF spaces measured on a macroscopic (organ level) and microscopic (tissue level) scale, we calculated the Spearman’s rank correlation between the normalized lateral ventricle volume and regional CSFF measurements.

## RESULTS

Figure 1 shows representative axial mid-brain WF maps obtained by FAST-T2 sequence from three subjects. Compared to the younger 22-year-old (Fig.1a) and 50-year-old (Fig.1b) subjects, the 73-year-old subject (Fig.1c) showed prominent structural brain changes, including cerebral atrophy, cortical loss, and ventricular dilation seen on the T1W image, as well as increased values in the brain parenchyma on the CSFF map.

**Figure 1.**
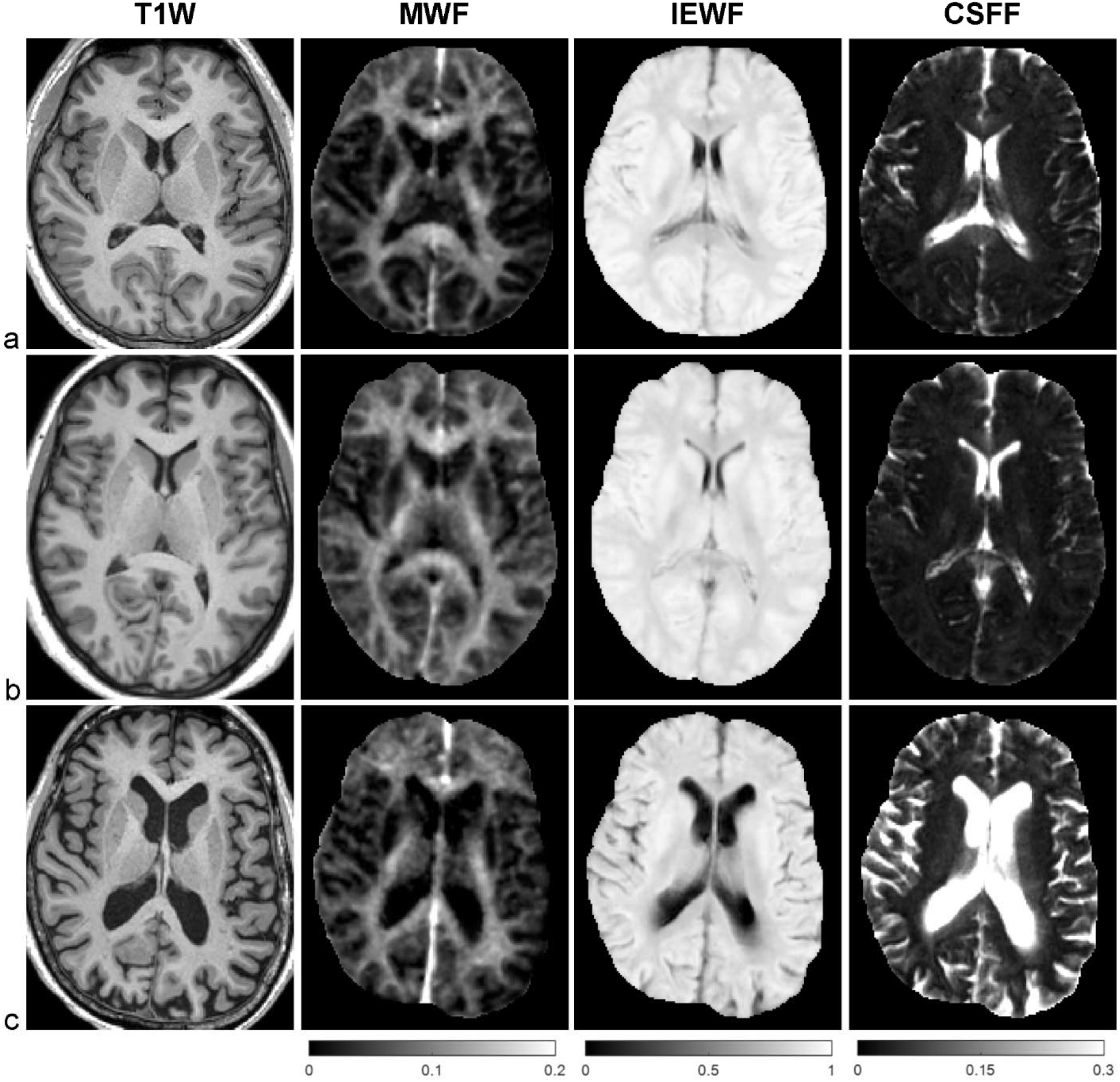
Representative axial mid-brain structural T1W MPRAGE image as well as corresponding MWF, IEWF, and CSFF maps derived from FAST-T2 data acquired from a) a 22-year-old male, b) a 50-year-old female, and c) a 73-year-old female CN volunteers. Note the marked increase in cerebral atrophy, cortical loss, and lateral ventricular volume on the T1W image, and higher parenchymal CSFF values in the latter subject compared to the younger subjects.

Figure 2 shows the scatter plots and corresponding second-order polynomial trend lines of regional MWF, IEWF and CSFF values versus age obtained in the cerebral WM, cerebral cortex, and cerebral deep GM. MWF is relatively stable and mainly decreases in the cerebral WM and deep GM of subjects aged 70 years and above, while IEWF generally decreases with age. In all three ROIs, mean CSFF values follow a quadratic trend and start to rise at approximately 50 years of age. Multiple linear regression analysis with FDR correction revealed a statistically significant quadratic relationship between age and regional CSFF measurement (QM model) for all three ROIs (all p-values < 0.001), after adjusting for sex and ROI volume. A statistically significant quadratic relationship with age was also found for MWF in the cerebral deep GM (p = 0.004) and IEWF in the cortex (p = 0.012). There was a statistically significant linear relationship between age and regional IEWF (LM model) in the cerebral WM (p = 0.006) and deep GM (p = 0.002).

Overall, the normalized cerebral WM and deep GM volumes were found to be stable up to approximately 50 years of age and decrease with age afterwards, accompanied by an increase in the size of the lateral ventricles (Fig.3). The normalized volume of the cerebral cortex shows a linear decreasing trend with age (Fig.3). Spearman’s rank correlation analysis showed a moderate correlation between the normalized lateral ventricle volume and the regional CSFF measurement obtained in the cerebral WM (ρ = 0.43, p < 0.001), cortex (ρ = 0.43, p < 0.001), and deep GM (ρ = 0.49, p < 0.001).

**Figure 2.**
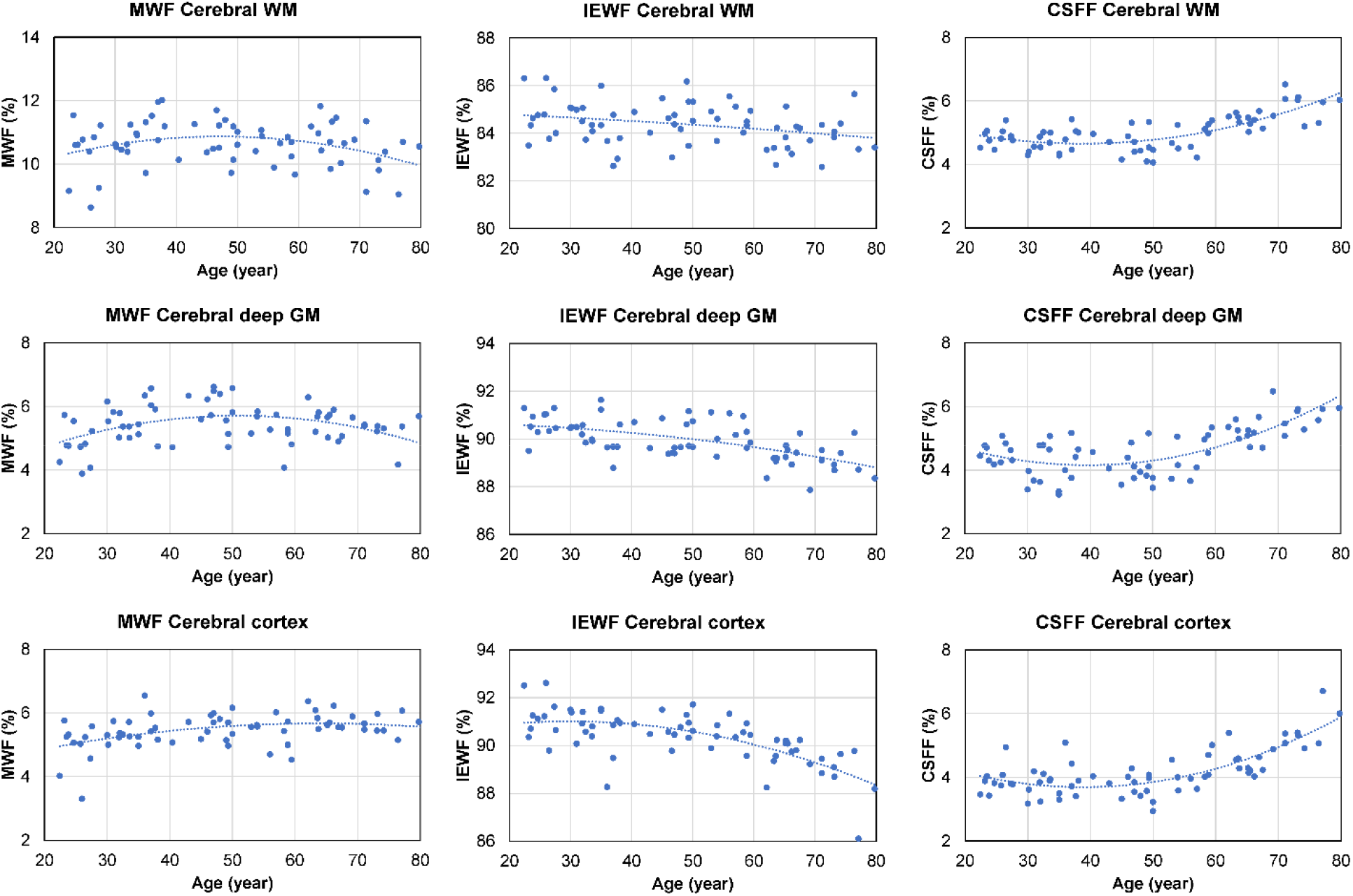
Scatter plots and trend lines (fitted using a second-order polynomial function) showing cross-sectional age-dependent MWF, IEWF and CSFF change measured in the cerebral WM, cortex, and deep GM of 66 CN subjects.

**Figure 3.**
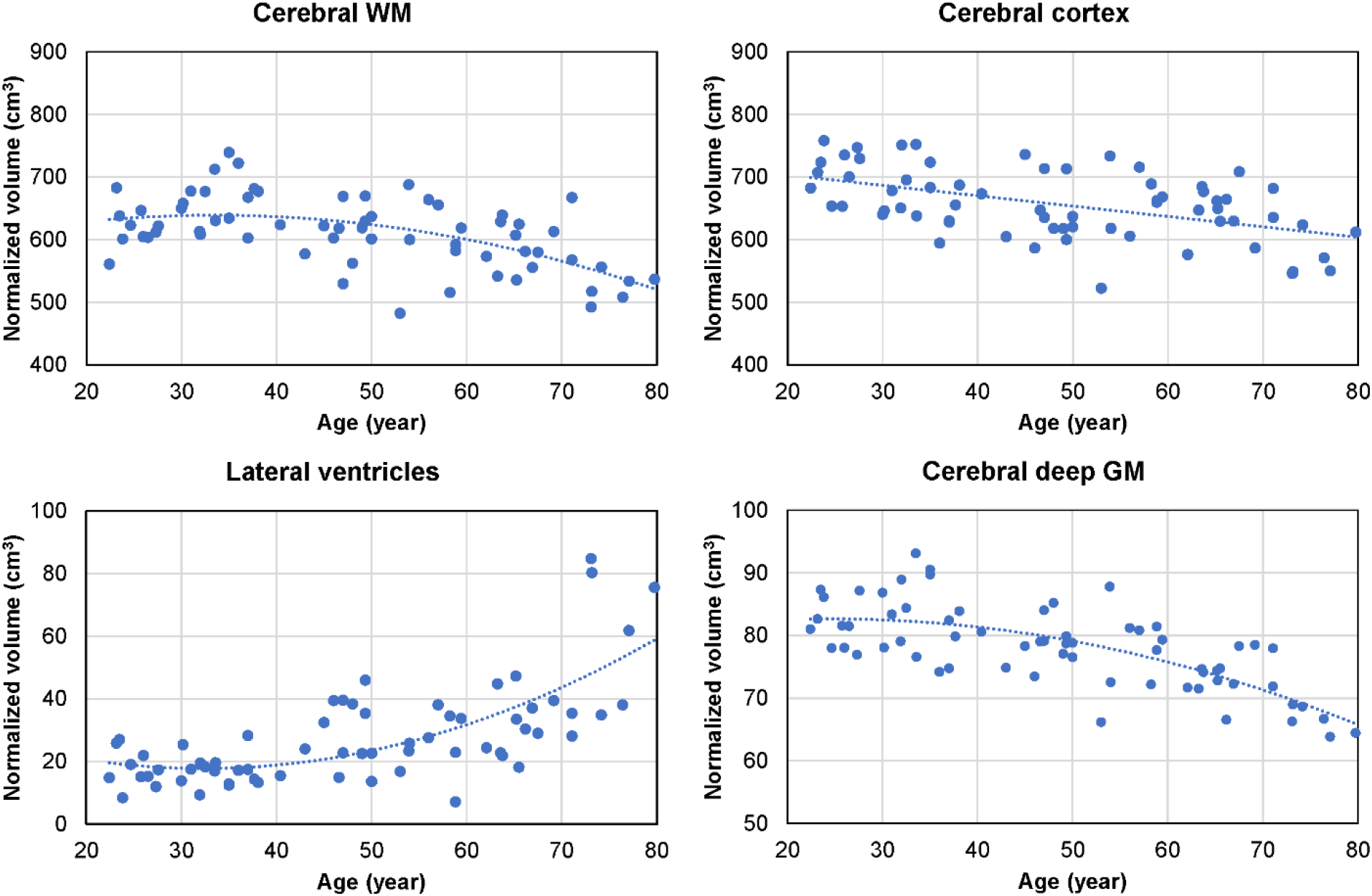
Cross-sectional brain volume data (normalized by the subject’s skull volume to account for difference in head size) obtained from 66 CN adults. The volume of the cerebral WM and subcortical deep GM shows a quadratic decreasing trend starting in the middle age (approximately at age 40). This is accompanied by a quadratic increase in the volume of the lateral ventricles, with notable enlargement observed in some subjects in the 70-80 age group. The volume of the cerebral cortex exhibits a gradual linear decreasing trend with age.

## DISCUSSION

In this cross-sectional MRI study of age-related changes in compartmental brain tissue water of CN subjects aged 20-80 years, we found strong evidence that parenchymal CSFF, a biomarker of the microscopic-scale CSF space, increases with age starting in the 6^th^ decade of life. After adjusting for sex and brain ROI volume, we found a highly significant quadratic relationship between age and regional CSFF in all three cerebral ROIs (WM, deep GM, and cortex). To the best of our knowledge, this finding has not been reported in the literature and may provide new insights into reorganization of brain tissue water and its association with CSF-driven glymphatic clearance. This is to be contrasted with the intra/extra-cellular water compartment, as measured by IEWF, that decreases linearly with age in the cerebral WM and deep GM and quadratically in the cortex. The myelin water component, expressed as MWF, is relatively stable with age and appears to follow a slightly inverted U-shape in the cerebral deep GM and WM, although a statistically significant quadratic relationship was found only in the deep GM. The different temporal patterns of the regional CSFF, IEWF and MWF change with age (Fig.2) likely reflect biological and physiological changes in an aging brain (Liu et al., 2017;Banks et al., 2021). Further studies are warranted to elucidate the biologic correlates of the observed WF changes and their roles in the pathogenesis and progression of age-related neurodegenerative diseases.

Among the three brain tissue water compartments studied in this work, the CSF component is often regarded as a confounder in brain MRI techniques, including functional MRI and proton MR spectroscopic studies (Horska et al., 2002), and has not been studied extensively in the context of aging. Qin et al performed single-compartment mapping of parenchymal CSF volume fraction (VCSF) using a T2-prepared 3D GRASE sequence in six healthy volunteers aged 32-50 years (Qin, 2011). They proposed to isolate the tissue CSF signal (T2 ∼ 2 s) from the signal of the other tissue water compartments (T2 < 120 ms) by acquiring multi-echo image data at long TEs > 600 ms. The data was fitted with a mono-exponential function to obtain the CSF signal at zero TE, which was then normalized by the ventricular pure CSF signal to obtain VCSF map. They reported mean VCSF of 8.9%, 11.4%, and 21.4% in the occipital, temporal, and frontal cortex, respectively, and VCSF of 0-3% in WM voxels. Assuming brain GM and WM water content of 83% and 70% (Whittall et al., 1997), respectively, and cortical CSFF of 3.8% and WM CSFF of 4.7% for the 30-50 years age group (Fig.2), we obtained a VCSF of 3.2% and 3.3% in the cerebral cortex and WM, respectively. The much higher cortical VCSF values reported by Qin et al compared to ours are most likely due to the larger partial volume effect in their image data (voxel size = 53.7 mm^3^ versus 7.8 mm^3^ in our study). Also, we should note that a conservative tissue mask was used in our ROI analysis (the distance between ROI voxels and CSF-filled brain reservoirs, including the ventricles and the subarachnoid space, is at least 1 mm in-plane and 5 mm through-plane), which further reduces partial voluming with the macroscopic CSF signal. In a more recent study (Canales-Rodriguez et al., 2021), Canales-Rodríguez et al applied a voxel-based analysis to three-compartment T2 relaxometry data obtained using a 3D GRASE sequence (Prasloski et al., 2012) in 145 healthy subjects aged 18-60 years. They found that parenchymal CSFF (termed “free and quasi-free water fraction” in their work) in general increases linearly with age, with a much larger slope observed in men than in women. Their reported brain CSFF values, ranging approximately 5-18%, are also higher than ours, which is likely due to the increased partial volume effect (voxel size = 42.9 mm^3^ versus 7.8 mm^3^ in our study) and differences in the data acquisition and post-processing methods. An important extension of our work over the study in (Canales-Rodriguez et al., 2021) is the inclusion of CN subjects in the 60-80 years age group and it is precisely in this group that a strong age effect on brain CSFF is demonstrated.

In this study, we observed a slightly inverted U-shape in the age-dependent MWF change in the cerebral deep GM and WM (Fig.3). A statistically significant quadratic relationship was found in the deep GM but not in the WM, possibly due to the limited sample size. While MWF is an established quantitative marker of myelin loss in demyelinating diseases such as MS (Mancini et al., 2020;van der Weijden et al., 2021), previous investigations in healthy individuals have reported inconsistent MWF trends with age, which could increase (Flynn et al., 2003;Lang et al., 2014;Canales-Rodriguez et al., 2021) or decrease (Faizy et al., 2018) linearly, remain stable (Billiet et al., 2015), or follow a quadratic inverted U-shape (Arshad et al., 2016;Papadaki et al., 2019;Bouhrara et al., 2020;Dvorak et al., 2021). The age-dependent quadratic behavior has also been observed for myelin-sensitive diffusion tensor imaging (DTI) indices such as fractional anisotropy or radial diffusivity (Lebel et al., 2012). As age-related changes in myelin sheath, axons, and myelin-producing oligodendrocytes in humans are complex and not well understood (Liu et al., 2017), further validations of the specificity of MWF and other myelin biomarkers to myelination in normal aging are needed.

The rise in parenchymal CSFF starting at approximately 50 years of age (Fig.2) is accompanied by the enlargement of the lateral ventricles (Fig.3), posing the question whether these two phenomena are related. Since the dilation of the lateral ventricles is a measure of cerebral atrophy (Nestor et al., 2008), we adjusted for the effect of age-dependent cerebral ROI volume on CSFF in the multiple linear regression model and found that age remained a statistically significant variable. We also found that the correlation between the lateral ventricle size and the regional CSFF measurements was statistically significant but not strong (Spearman’s correlation < 0.5). The correlation was notably poor for subjects in the 70-79 years age group, with half of the group having much larger lateral ventricles (Fig.3) yet similar regional CSFF values compared to the other half. These results suggest that the macroscopic (organ level) and microscopic (tissue level) CSF spaces are related, but CSFF is a tissue marker independent from the observable macroscopic brain atrophy and ventricular dilation. A possible interpretation of CSFF increase with age is the dilation of the microscopic PVS due to reduced water transport (Ford et al., 2022) and impaired glymphatic waste clearance in the aging brain (Nedergaard, 2013;Wardlaw et al., 2020). PVS enlargement is conventionally detected and quantified on the T2W image (Ramirez et al., 2016;Ballerini et al., 2018). A recent cross-sectional study of 1789 healthy subjects aged 8-100 years revealed a biphasic PVS volume change with age in the basal ganglia and the WM, slightly decreasing until the mid-30s and increasing afterwards (Kim et al., 2022), which is very similar to the trends seen in our study (Fig.2). However, unlike CSFF, there was a considerable variation in PVS volume fraction, especially in the older subjects (Kim et al., 2022). Since this approach is not capable of detecting PVS much smaller than the voxel size (approximately 1 mm), future works are needed to study the association between the T2W-visible PVS load and CSFF in the same cohort.

Several studies have reported a decreasing trend with age of IEWF in various brain regions (Billiet et al., 2015;Canales-Rodriguez et al., 2021), which supports our findings. Between the age of 20 and 80 years, we found the reduction in IEWF is about 1% in WM and 2% in GM (in absolute terms), which is accompanied by a similar increase in CSFF. Since the intra/extra-cellular water compartment is the dominant source of water in the brain tissue (IEWF ∼ 90% in GM and 85% in WM), a decrease in this water compartment alone does not fully explain a corresponding increase in the tissue CSF water. As an illustrative example, assume the water content of GM is 1 g/mL, which consists of 0.05 g/mL myelin water, 0.90 g/mL intra/extra-cellular water, and 0.05 g/mL CSF. A loss of water in the intra/extra-cellular compartment alone (from 0.90 g/mL to 0.73 g/mL) will result in a 2.0% decrease in IEWF, but only a 1.0% increase in CSFF. Therefore, the observed CSFF increase of 2% indicates that there is an increase in the CSF compartment. To ascertain the true magnitude of change in these water compartments, a voxel-wise mapping of the brain tissue total water content (Neeb et al., 2006;Meyers et al., 2017;Nguyen et al., 2017) is needed and will be considered in our future work.

The present study has several limitations. First, our sample size is relatively small, which restricts the number of ROIs upon which inference can be performed in the statistical analysis. Second, due to the scan time constraint, we did not measure the tissue water content, which makes the interpretation of the changes in the compartmental WF, a relative measure, somewhat challenging. Third, current multi-component T2 relaxometry techniques cannot differentiate intra-cellular and interstitial water compartments within a voxel. In our future work, we will consider combining FAST-T2 with diffusion imaging approaches such as neurite orientation dispersion and density imaging (NODDI) (Zhang et al., 2012) and free-water DTI (Pasternak et al., 2009) which are capable of separating these water compartments. Fourth, the FAST-T2 imaging slice is relatively thick (5 mm), which necessitates ROI mask erosion to mitigate the partial volume effect. Finally, we made the assumption that CSFF mainly captures the contribution from CSF water, which is reasonable based on the current understanding of the water T2 spectrum in the brain tissue (MacKay et al., 2006). Further independent validations, by histology or other imaging methods, are required to confirm the findings in our work.

In conclusion, brain tissue water residing in different water compartments shows complex changing patterns with age over the adult lifespan of 20-80 years. Parenchymal CSFF, a biomarker of microscopic-scale CSF-like water, shows a quadratic increase in both GM and WM, starting approximately at the age of 50.

## Conflict of Interest

The authors declare that the research was conducted in the absence of any commercial or financial relationships that could be construed as a potential conflict of interest.

## Availability of the data

Data available upon reasonable request given the need for a formal data sharing agreement between the authors’ and the requesting researchers’ institutions.

## Acknowledgements

The authors thank Dr. Silky Pahlajani and Emily B. Tanzi for their help with subject recruitment and characterization.

## Notes

### Competing Interest Statement

The authors have declared no competing interest.

